# STAMPede: A Precision Oncology Catalogue of Solid Tumors

**DOI:** 10.1101/2020.09.18.290635

**Authors:** Jessica W. Chen, Haik Kalantarian, Christian A. Kunder, James L. Zehnder, Henning Stehr, Helio A. Costa

## Abstract

Molecular profiling of tumor specimens is a key contributor in the application of precision medicine toward patient care in oncology. The presence of genetic mutational data may provide insight into the etiology of the cancer and inform about the available therapeutic options for patients diagnosed with various cancer types. However, the ability to centrally organize, structure, and visualize this genetic data can be hindered by the lack of necessary computational infrastructure. Here we present a somatic tumor data visualization portal titled STAMPede that utilizes tumor sequencing data from an in-house solid tumor oncology sequencing panel at Stanford Health Care. STAMPede is intended to provide Stanford healthcare providers and clinical researchers an easy to navigate web-based portal to query and display gene-, variant-, and cancer-level summary statistics.

## INTRODUCTION

The application of next-generation sequencing (NGS) to profile genetic alterations in tumor specimens of patients has impacted the therapeutic treatment and care management of patients, specifically, enabling the discovery of diagnostic, prognostic, and predictive biomarkers^1^. With the deluge of data generated by routine somatic tumor sequencing are gaps in the essential infrastructure to permit healthcare providers and clinical researchers to systematically organize and visualize these sequencing results. Oncology-based genetic databases offer opportunities to transform the landscape of biomedical research and clinical development. In alignment with ongoing precision medicine efforts^2,3^, analysis of aggregated genetic data creates a platform that offers the opportunity to accelerate the identification of novel biomarker-based indications for which existing pharmaceutical drugs may be found to be efficacious, and thereby, address therapeutic areas of unmet need. There currently does not exist a clinical visualization platform at Stanford Health Care to disseminate summary-level statistics of the genetic data from the STanford Actionable Mutation Panel (STAMP) family of assays, a targeted next-generation sequencing assay platform for tumor biopsy specimens. In our study, we developed an in-house browser that leverages the existing structure of the somatic genotyping data to provide a web-based interactive tool that enables exploration of the relationship between cancer type and somatic alterations. This resource provides a foundation for future prospective and retrospective exploratory analyses and provides high-level data to healthcare providers regarding the spectra of genetic alterations that exist within their patient populations.

## MATERIALS AND METHODS

### Source of solid tumor molecular data

The solid tumor biopsies from patients described in this study were obtained from formalin-fixed paraffin-embedded tissue blocks from Stanford Health Care as part of routine patient care in oncology. Quality controls measures have been previously described^4^.

Somatic molecular testing was performed on solid tumor biopsies using the NGS-based STAMP assay offered by the Stanford Molecular Pathology laboratory, which is a CLIA-certified and CAP-accredited clinical diagnostic laboratory [https://stanfordlab.com/content/stanfordlab/en/molecular-pathology/molecular-genetic-pathology.html/]. The assay identifies somatic alterations (single nucleotide variations [SNVs], insertions/deletions [Indels], copy number variations [CNVs], and gene fusions) in 130 genes that have been implicated in cancer. For this analysis, a unique patient is defined by a unique test order identifier.

### Implementation

STAMPede data visualizations are implemented in R using the Plotly framework, an interactive browser-based graphic library for R (Plotly, Montreal, Canada). The source code is available in the Github repository: https://github.com/SHCMolPathLab/Jessica/tree/master/STAMPEDE_Visualizations.

### Web Portal Implementation

The STAMPede web portal provides a gateway for the visualization of summary-level statistics of the STAMP database and is implemented using HTML/Javascript/CSS. The portal is hosted on Stanford’s Andrew File System service and uses CSV files linking to the Plotly data visualizations as the primary datastore. The web page restricts access via a whitelist to Stanford clinicians and clinical researchers using an Apache .htaccess file.

## RESULTS

### Browser features

#### STAMPede Homepage

The homepage presents summary-level statistics of the entire STAMP database (**Figure 1**). The occurrence frequency of all genes on the STAMP panel (cumulative and filtered for the fusion genes), and the primary site of the tumors in the patient population, are visualized in histograms. In addition, the top occurring (in terms of quantity) genes, fusion gene pairs, variants, and primary tumor sites, are visualized in histograms. Specifically, those with frequencies equal to or greater than that of the 20th top gene, fusion gene pair, variant, or primary tumor site, respectively, will be visualized. Moreover, the distribution of variant types (i.e. SNV/Indels, CNV amplifications, and fusions) is visualized in a histogram. For SNV/Indels, additional stratification is provided through the variant type classifications of SNV, frameshift Indel, in-frame Indel, and the pathogenicity classifications of pathogenic variants and VUS. Lastly, the gender and age demographics of the patient population is visualized in a histogram.

**FIGURE 1.**
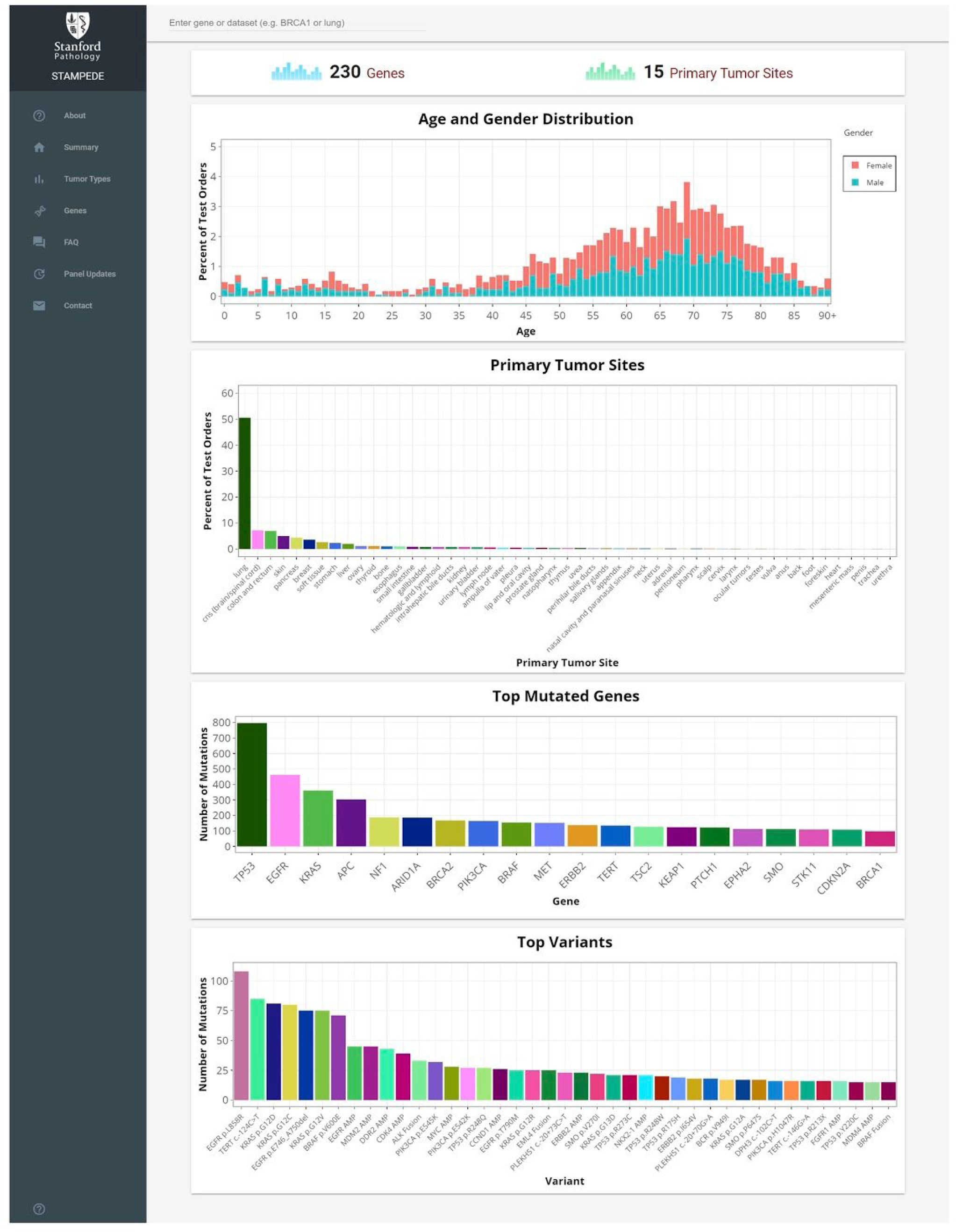
STAMPede homepage with summary level visualizations of the entire STAMP database. The browser enables summary-level examination of the relationship between phenotype (i.e. tumor type) and genotype (i.e. identified somatic mutations -SNVs, Indels, CNVs, and fusions). The STAMPede homepage summarizes all mutations in the STAMP database and displays the occurrence frequency of all genes on the STAMP panel (cumulative and filtered for the fusion genes), and the primary site of the tumors; the top occurring genes, fusion gene pairs, variants, and primary tumor sites; the variant type distribution; and the gender and age demographics of the patient population.

#### Gene stratification

The gene page presents summary-level statistics stratified by gene (**Figure 2**). For each gene, the top fusion gene pair, variants, and primary tumor sites are visualized in histograms. For the top occurring fusion pairs for a gene of interest, if all the fusion pairs occurs only once, the first 20 fusion pairs when sorted alphabetically will be visualized; if at least one fusion pair occurs more than once and there are less than 20 fusion pairs, all the fusion pairs will be visualized; if at least one fusion pair occurs more than once and there are more than 20 fusion pairs, the fusion pairs with frequencies equal to or greater than the 20th top fusion pair will be visualized. For the top occurring variants for a gene of interest, the decision tree logic applied is the same as that of the top occurring fusion pairs. For the top occurring primary tumor sites for a gene of interest, if there are less than 20 sites, all the sites will be visualized; if there are more than 20 sites, the sites with frequencies equal to or greater than the 20th top site will be visualized and all sites that occur only once will be removed. In addition, the variant type distribution is visualized in a histogram and the frequency of each SNVs along the 2-dimensional sequence of the gene is visualized in a lollipop diagram with the motif domains annotated.

**FIGURE 2.**
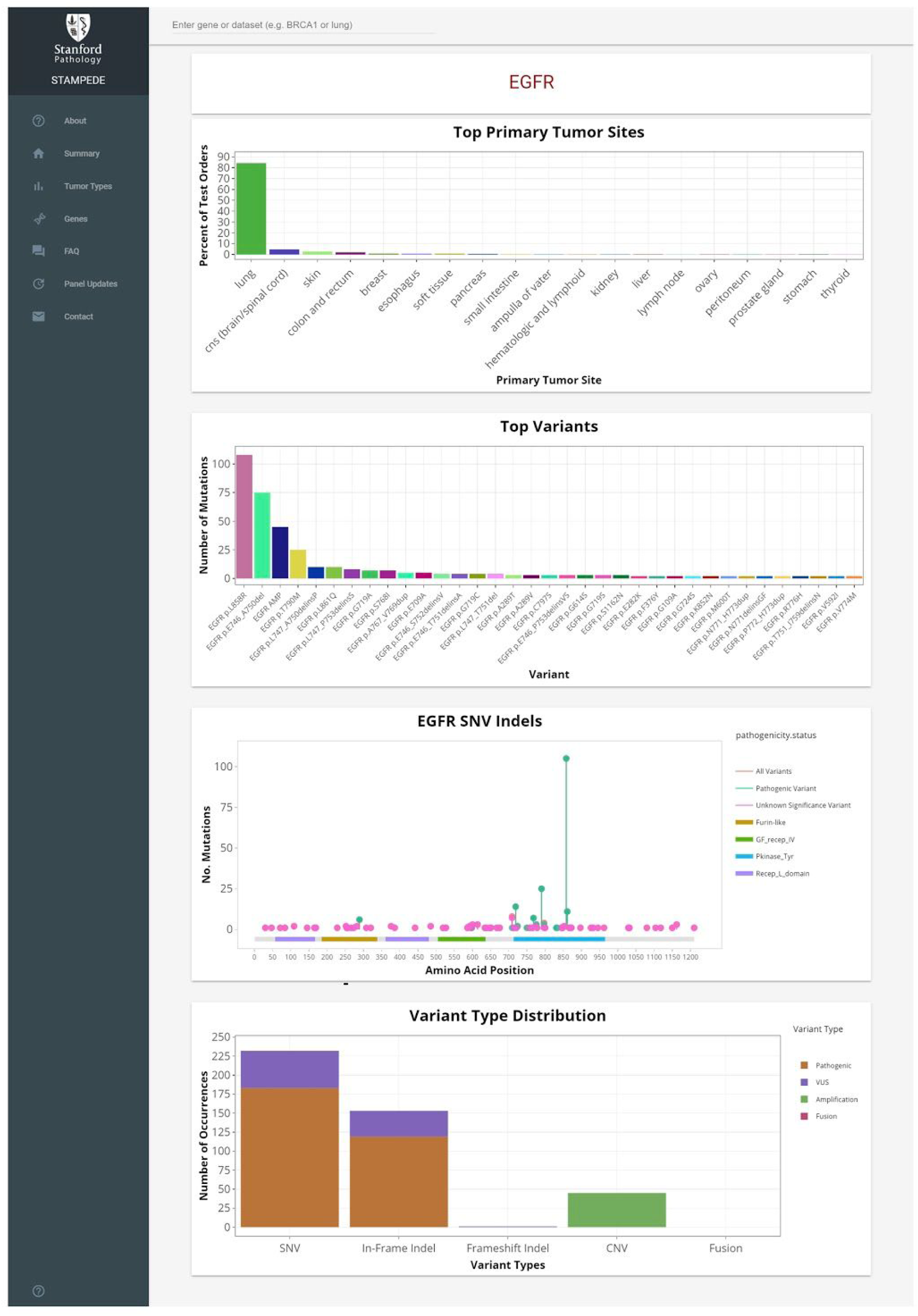
Visualization of STAMP database stratified by gene (i.e. EGFR). The visualizations summarize all gene-specific (i.e. EGFR) mutations in the STAMP database and displays the top fusion gene pair, variants, and primary tumor sites; the variant type distribution; and the frequency of each SNVs along the 2-dimensional sequence of the gene.

#### Tumor type stratification

The tumor type page presents summary-level statistics stratified by primary tumor site (**Figure 3**). For each primary tumor site, the top fusion gene pairs, genes, and variants are visualized as histograms. For the top occurring fusion pairs and variants for a primary tumor site of interest, the decision tree logic applied as the same as that described for the gene stratification page. For the top occurring genes for a primary tumor site of interest, if there are less than 20 genes, all the genes will be visualized; if there are more than 20 genes, the genes with frequencies equal to or greater than the 20th top gene will be visualized and all genes that occur only once will be removed. In addition, the gender and age demographics of the patient population for the primary tumor site of interest is visualized in a histogram.

**FIGURE 3.**
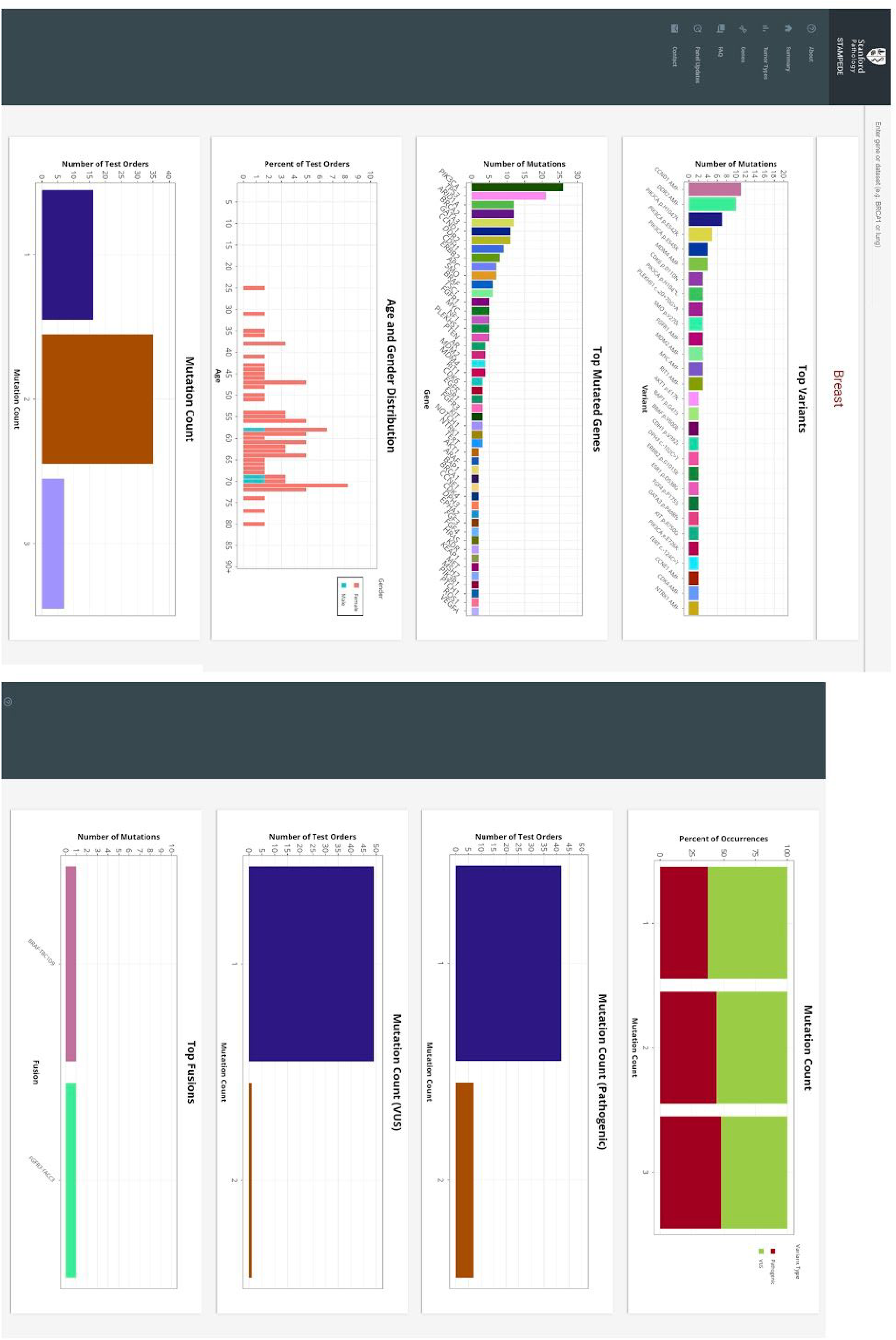
Visualization of the STAMP database stratified by tumor type (i.e. breast cancer). The visualizations summarize all tumor-specific (i.e. breast cancer) diagnoses in the STAMP database and displays the top fusion gene pairs, genes, and variants; and the gender and age demographics of the patient population.

## DISCUSSION

The STAMPede catalogue aims to provide a resource to clinicians and scientific researchers at Stanford Cancer Institute to assist clinical decision making and aid in clinical research projects. Currently available precision medicine knowledge bases include OncoKB^5^, CIViC^6^, MyCancerGenome [https://www.mycancergenome.org], and the Jackson Laboratory Clinical Knowledgebase^7^, among others. While the genes and testing panels used to generate interpretive comments across these resources are not the same^8^, there is an overlap of a majority of actionable mutations and general consensus around their clinical interpretation. Caveats include differences in the bioinformatic pipelines used for processing of the raw files, which may result in differences in the criteria for calling a mutation and inclusion within these resources.

Collectively, these catalogues shed light on the genomic landscape of somatic alterations in oncology as the data is derived from a multitude of large-scale institution-wide sequencing efforts. As precision medicine efforts in the drug development space are headed towards more targeted therapies^9^ and the development of basket trials such as the NCI-MATCH trials^10^, these catalogues provide evidence-based support for testing FDA-approved drugs in new indications. For example, biomarker prevalence derived from the catalogues may inform forecasting of the unmet medical need in the oncology community and thereby help guide clinical development plans.

## CONCLUSION

Our study describes an interactive catalogue that aggregates the genetic alterations identified by the STAMP NGS assay and generates summary-level visualizations of the data as a platform to support precision medicine efforts in oncology. The gene-, variant- and cancer-level data presented in STAMPede enables both healthcare providers and researchers to better understand the genetic underpinning of the patient populations they directly serve. Additionally, this resource will enable retrospective exploratory analyses that combine electronic medical records and genomic sequencing data to identify putative prognostic and predictive biomarkers.

## AUTHOR CONTRIBUTIONS

HAC conceived and initiated the project. JWC, HK, HS, and HAC designed and performed the study. HAC, HS, and JLZ supervised the work. JWC, HK, and HAC wrote and edited the manuscript.

## AVAILABILITY

STAMPede browsing capabilities are available for access controlled healthcare providers and clinical researchers at stampede.stanford.edu.

## DISCLOSURE

J.W.C. is currently a full-time employee at Genentech, Inc. and holds stocks in Roche Holding AG. None of these entities played a role in the design, execution, interpretation, or presentation of this study.

